# TAS2R receptor response helps predict new antimicrobial molecules for the 21^st^ century

**DOI:** 10.1101/2022.10.25.513703

**Authors:** S Sambu

**Affiliations:** Advanced Computational Sciences Department, Bartanel Discovery, Turnhoutsebaan 139 A, Antwerp 2140, Belgium

**Keywords:** Machine Learning, antimicrobial, artificial intelligence, bitterness, QSAR, TAS2R

## Abstract

Artificial intelligence (AI) requires the provision of learnable data to successfully deliver requisite prediction power. In this article, it is demonstrable that standard physico-chemical parameters, while useful, were insufficient for development of powerful antimicrobial prediction algorithms. Initial models that focussed solely on the values extractable from the knowledge on the electrotopological, structural, constitutional descriptors did not meet the acceptance criteria for classifying antimicrobial activity. In contrast, efforts to conceptually define the diametric opposite of an antimicrobial compound helped to advance the category description into a learnable trait. Interestingly, the inclusion of ligand-receptor information using the ability of the molecules to stimulate transmembrane TAS2R receptor helped to increase the ability to distinguish antimicrobial molecules from the inactive ones. This novel approach to the development of AI models has allowed the development of models for the design and selection of newer, more powerful antimicrobial agents. This is especially valuable in an age where antimicrobial resistance could be ruinous to modern health systems.

## 1 Introduction

Quantitative structure-activity relationship (QSAR) models are essential tools for drug development because they can be used to assess large chemical libraries in the hunt for new antimicrobials. However, for chemical antimicrobials, one of the strongest challenges to this approach is the rarity of powerful antimicrobials in the vast chemical spaces that are the presumed relevant chemical spaces. If they were more abundant, the hunt for antimicrobials would be sooner met. This rarity means that structure and activity descriptors are often insufficient to describe the variability in the antimicrobial datasets.

Aside from the insufficiency of descriptors for the chemical space, the modelling dataset is often severely imbalanced. That is, of the two initial classes for categorizing the chemistries, one will be highly populated (the inactive class) while the other will be significantly diminished (the active class). This means that the problem is compounded since the many descriptors that require parametrization will now have to be learnt only after oversampling for the minority class whilst undersampling the majority class. Computing efficiencies and data restrictions prevent proper statistical matching during these popular balancing procedures. This learning method will result in less-than-stellar computing efficiencies and worse model metrics(1). Additionally, during real-world usage, the model will run the risk of encountering unfamiliar chemical spaces especially in the assignment of “*active*” antimicrobials.

Given the difficulty in the learning challenge posed by severe class imbalances, it may help to use descriptors that are sensitive and are mechanistically sound. Parameters that quantify factors underlying the molecular movement and interaction might be helpful in identify molecules that easily reach systemic circulation. However, there is a need to include rules that examine the ability to engage in flexible (ligand-receptor) binding as might be found in TAS2R receptors(2,3) well known for their polymorphism resulting in a “real life” variable shape and variform virtualized sensor with which to test potential target molecules.

A variform test to binding ability may be a more useful descriptor for antimicrobial activity. Effectively, a ligand’s psychophysical response to a candidate molecule encodes a molecular cypher whose contents relate a molecular key to physiologic aspects including toxicity and therapeutic effects(4–6). Hence, TAS2R ligands may contain significantly greater subsets of molecules with antimicrobial effects which subsets can subsequently be evaluated for safety/toxicity signals. A measure of TAS2R-triggering capacity of the molecule will provide a mechanistic flavour within the AI model and may enable the algorithm to better bridge across training-versus-test set while eliminating the need for a universal molecular embedding challenge.

Historical medical anthropological studies may also provide evidence of the TAS2R receptors as evolutionary molecular cyphers since bitter substances would elicit a learnable aversive response unless a therapeutic effect presented a large enough benefit to overcome the aversive barrier (7). Additionally, the ability to titrate sub-toxic levels against therapeutic thresholds of a candidate TAS2R trigger species could explain the association between bitter cyanogens and observable nonclinical parasitaemia in West African communities (7). These pieces of evidence motivate the inclusion of TAS2R response in the drug design workflow.

Overall, the use of antimicrobial TAS2R ligands may provide us with more powerful dual-action antimicrobials. TAS2R ligands are known to trigger the release of mammalian antimicrobial compounds including peptides that aid in the resolution of microbial infection(8,9). Adding this effect to the direct microbicidal activity of the ligand will enhance the therapeutic effect of the molecule. Given that the TAS2R immune-activating antimicrobial effect works at micron-level thresholds(6), the effect of the treatment will also demonstrate an increased overall efficacy while maintaining pre-extant safety margins against potential toxicity.

The immune-activating properties of bitter compounds have been identified in bacterial quorum sensing molecules(10,11). This implies the existence of bitterness sensing receptors in the bacteria and the mammalian cells likely as part of a long-running evolutionary arms race. It is logical to conclude that a TAS2R psychophysical predictor variable(s) could help map immune activating chemistries onto an antimicrobial predicted variable.

Computationally, the psychophysical TAS2R response vector may help to embed the chemistry with the right biological projection. This is in contrast to numerous attempts at development of a universal molecular embedder. Typical approaches that attempt to generate standard fingerprints (FP) run into intractable challenges such as variability in the FP length(12), limitations in the representational scope of the embedder(13) and increasing computing inefficiencies with increasing molecular complexity(13). Using TAS2R response encoders helps alleviate these challenges all in a single bio-endogenic vectoral representation.

The dual-mechanism (immune trigger and antimicrobial activity) approaches may lessen the risk for antimicrobial resistance but they do not eliminate antimicrobial resistance. The requirement to regulate usage in both human and veterinary medicine is necessary to ensure durability of any next generation antimicrobial drug’s efficacy. Additionally, the exploration of genetically primed phages to target known microbes at a subspecies level via personalized medicine may also help to increase the breadth of the medical armoury against antimicrobial resistance(14).

## 2 Materials and Methods

### 2.1 Data Collection

Data was downloaded from an antimicrobial inhibition assay data deposited on PubChem(15) as part of a screening assay by the Southern Research Molecular Libraries Screening Centre (SRMLSC). The assay uses E. coli *BW25113 ΔtolC: Kan* as the test organism. *E. Coli* were exposed to the test compounds at fixed 30 µM dose grown in 384-well plates with inhibition readings obtained from optical density measurements at 615 nm on Envision® microplate reader(15). Samples were read at the same concentration(15). The dataset contains 65,120 compounds with 2.18% being active and 97.82% being inactive. Only molecules with significantly high antimicrobial activity (> 65 % inhibition) were considered for the active class; in like manner, only molecules showing negligible antimicrobial activity (< 35% inhibition) were included in the inactive class.

### 2.2 Model Development

#### SMILES Generation

Molecular SMILES were generated from chemical identifiers (CID) using a PubChem Database FTP service and then checked for consistency.

#### Variable Transformation

Molecular SMILES were read into R from an “SMI” file. The SMI object was used to generate parameters including octanol-water coefficients using atomic methods, atom molar refractivity (AMR), tallies for acid groups, base groups and rotational bonds, Numbers of Lipinski Failures and molecular weight. A bitterness index was generated using a primarily structure-based bitterness model(4).

#### Model Development

Using both the CDK descriptors and the bitterness index, a number of modelling approaches were attempted alongside grid-based hyperparameter optimization. These included distributed random forests (DRF), gradient boosted models (GBM) and stacked model ensembles. The stopping metric for model development was maximizing the area-under-curve on the precision-recall graph (AUCPR) for the validation set. In a later development, incoming data were statistically matched using the *MatchIt* package(16) using the number of Lipinski ‘s Rules met as a matching factor. The matched data were subsequently used in a model building exercised as described in this section.

## 3 Results and Discussion

### 3.1 Model outcomes meet standard quality criteria

The model performance met the intended success criteria given the high proportions of True Positives and True Negatives as shown in Table I (main diagonal values). Off-diagonal values (False Positives and False Negatives) are a significant minority bringing in an overall error rate of 9% (bottom right corner).

**Table I:** The confusion matrix summarizing the validation outcomes for the GBM model

|  | ACTUAL |  |  |  |
| --- | --- | --- | --- | --- |
| PREDICTED |  | Active | Inactive | Error |
|  | Active | 94% | 6% | 6% |
|  | Inactive | 12% | 88% | 12% |
|  | Totals | 58% | 42% | 9% |

### 3.2 Descriptor contributions to the model development

Molecular structure was used to generate a range of mechanistic and binding-related structural factors to build the prediction model. These factors are divisible into these main areas: mass, Hydrophobicity, electro-topological, binding and combinatorial representations. A summary of the descriptor choice is provided in Table II.

**Table II:** A representative summary of high variance features that contribute significant information to the model.

| Descriptor | Function | Contribution | Rationale |
| --- | --- | --- | --- |
| Mass | Molecular Weight | 10% | Diffusion rate across biomembranes & cytosol |
| Hydrophobicity | Multiple | 40% | Partitioning across biomembranes |
| Electro-topological | Polarizability | 10% | Charge separation in response to an external electric field: may influence coulombic interactions with cellular environment |
| Binding Degrees of Freedom (D.F.) | Multiple features | 20% | Solvation and motion free energies determined by acid/base pairs & rotatable bonds |
| Combinatorial | Multiple Features | 20% | Drug design rules and psychophysical response indices |

### 3.3 Analysis of variable importance underscores the value of the Bitterness Index

In Figure 1, the variable importance plot shows the most influential variables in the model. There are polarizabilities (*AMR*), partitioning indices (*ALogp2, ALogP, XLogP*), molar mass (*MW*) and binding/interactivity DoF (*nRotB & nBase*). These are mechanistic in nature driving the activity of each member. Polarizability refers to the impact of an electric field on the separation of charges within the molecule as might be displayed during electric separation (barrier effects) by the cell membrane to electrostatically exclude negatively charged molecules. Partitioning indices also affect diffusion across the cell membrane driven by the hydrophilic-lipophilic balance. Molar mass affects molecular diffusion directly with larger molecules moving slower than small ones across the biomembrane. Lastly, the ability for the molecule to attain preferrable solvation and motion free energies during the compartmental transition across the biomembrane is driven by molecular degrees of freedom (DF) for the biomembrane transition. These latter measures are composable into acid-base pairing and sub-molecular rotary motion. The bitterness index (*bi*) stands out in this non-stacked model, it being a psychophysical measure accounting for the ability of the molecule to trigger a TAS2R event. TAS2R are a family of known transmembrane receptors expressed extra-orally; TAS2R-activating molecules are known to have bacteriostatic and immunologic trigger effects(8,17).

**Figure 1:**
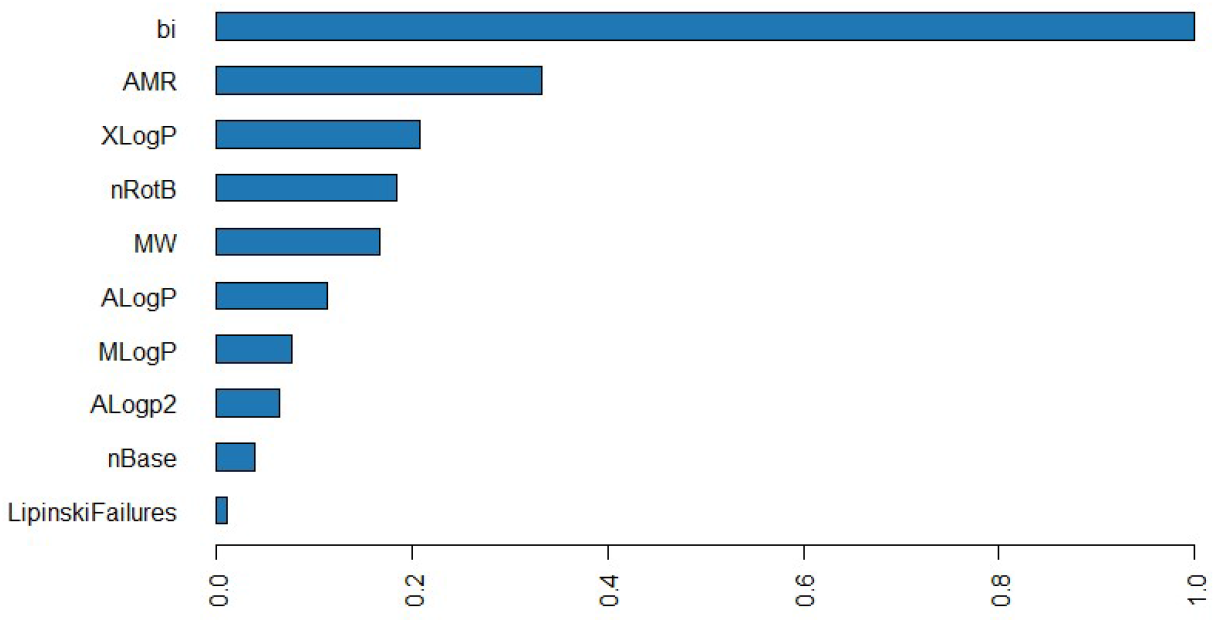
A variable importance plot showing the most influential variables in the DRF; *Validation metrics: AUCPR/AUC (>=0*.*95); Mean Error/Class (0*.*1)*

The whole-model contributions in a variable importance plot do not indicate the row-wise and directional contributions for each factor; for the mean marginal, combinatorial, row-wise contributions (in addition to the global structural understanding), SHAP values are required(18). In Figure 2, Shapley additive explanations (SHAP) values in the validation set show that the bitterness index (*bi*) is the most significant (topmost factor) and has negative contributions delineated from positive contributions. Additionally, high scalar *bi* values (red) are distinguished from the mid and low-level values of the bitterness index. Molecular weight (*MW*) also shows a similar separation between low and high MW species. AMR demonstrates clear centroids with molecules with low polarizabilities having a higher contribution to antimicrobial activity. Hydrophobicity measures demonstrate a head-tail morphology in their contributions between their column-specific normalizations typical of atomistic contribution models for partitioning. Acid-base counts are linked mechanistically according to donation/reception of hydrogen ions during molecular interaction with only very high values demonstrating robust SHAP value contributions. Lipinski Failures (LF) have very little contributions which may point to the potential confoundedness of the variable (see section 3.4.3). LF may rightly affect the antimicrobial activity from a design/selection perspective and similarly are affected by the independent factors such as molecular weight and binding/interactivity parameters. Hence, a de-confounding process may help to improve the model’s determinacy.

**Figure 2:**
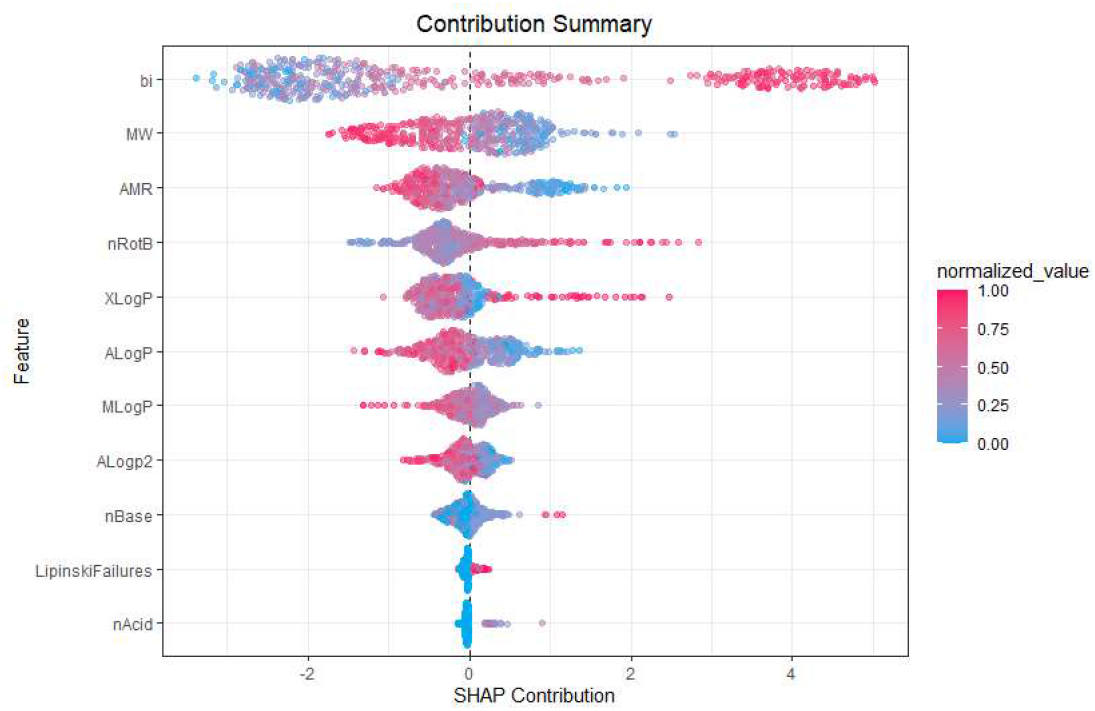
SHAP values applied against the validation set show that bitterness index (*bi*) has clear statistical centroids with non-zero Shapley value contributions. *Validation metrics: AUCPR/AUC (>=0*.*99); Mean Error/Class (0*.*04)*

#### 3.3.1 The bitterness index as a water-shed differentiating variable

By plotting the bitterness index against the antimicrobial classes, as shown in Figure 3, it is demonstrable that the feature separates neatly. Active compounds have significantly lower mean bitterness indices (centreline connecting the two notches) overall than inactive ones. There are exceptions to this general rule (shown in the red “cross” outliers) which may indicate that the classification challenge is a multivariate one.

**Figure 3:**
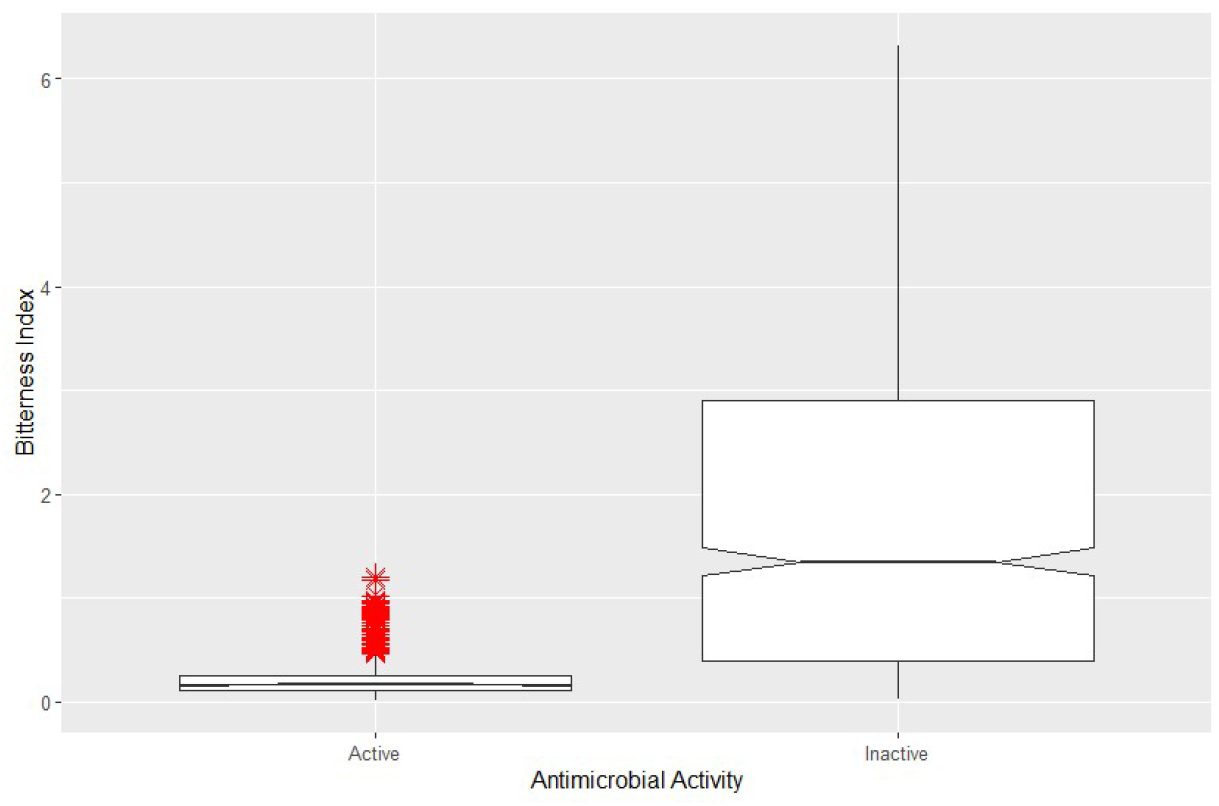
A notched box plot showing the ability of the bitterness index to distinguish between the two categories for antimicrobial activity (*p < 0*.*05; signified by the notch*). Antimicrobial activity can have a broad continuous range which requires a feature like bitterness to separate the two classes to their right statistical centres.

A similar examination can be rendered for mean molecular weight (centreline connecting the two notches) which also appears to separate the two classes for antimicrobial activity well as shown in Figure 4. Although this separation is not as exacting as bitterness, it is nonetheless indicative of the value of molecular weight within a classification model for antimicrobial activity.

**Figure 4:**
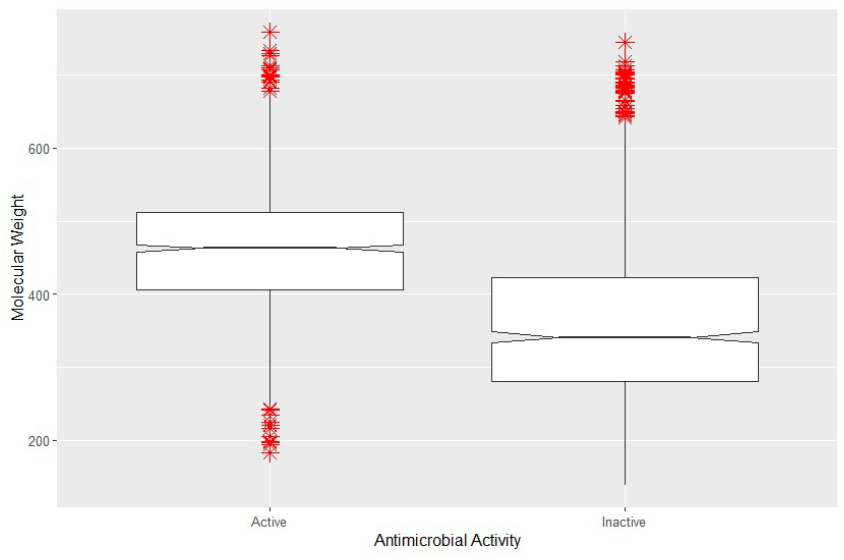
A notched box plot showing the ability of the molecular weight to distinguish between the two categories for antimicrobial activity. The notches indicate there is a significant difference in the means of the two groups (*p < 0*.*05*).

#### 3.3.2 Eliminating the bitterness index impoverishes model performance

Additional testing to examine whether the versions of models that lowered the importance of the bitterness index could be drawing out simpler models was warranted. Hence, by eliminating the bitterness index, one could test whether the resultant model quality was degraded or remained competitive with earlier versions containing the bitterness index. Overall, from Table III, the bitterness index is rather useful since the model without the bitterness index suffered a 67% reduction (overall score) in quality especially as measured by the maximum mean error and the max absolute Matthew’s correlation coefficient (MCC)

**Table III:**
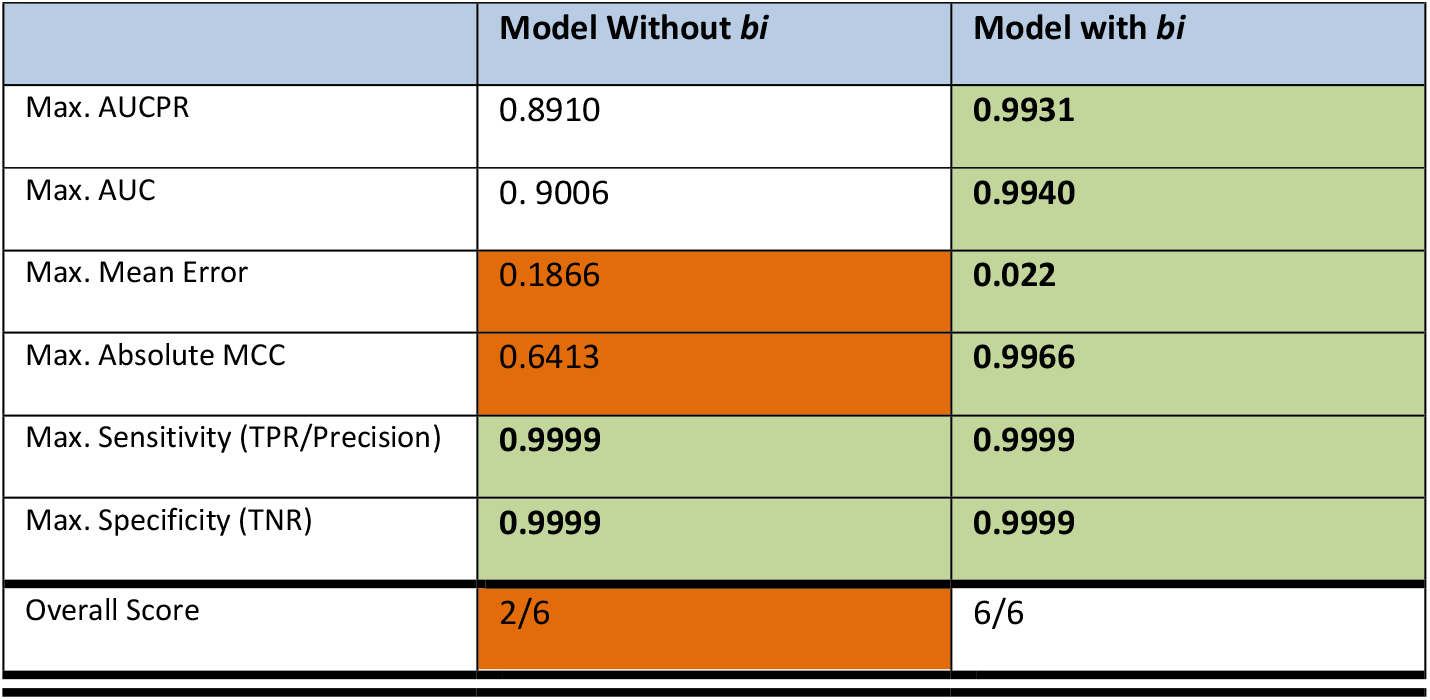
Performance comparison (maximum performance conditioned on respective thresholds) of top performing models with and without the bitterness index (*bi)* shows that the *bi* vector is critical for attain quality model parameters. Noticeably, given the dataset is imbalanced, the MCC (Matthew’s Correlation coefficient), which accounts for all true-false/positive-negative rates shows that the bitterness index is essential to achieving a reasonable model. The maximum mean error (MME) also shows that without the bitterness index, the model suffers from approximately 10x increase in the observed MME. The overall score shows that only 33% (2/6) of the model quality parameters are comparable to the model with the bitterness index.

### 3.4 An analysis of model development

#### 3.4.1 Contrastive evaluation of cooperative contributions recapitulates feature selection

To examine the magnified row-level contribution impact, it may help to select two contrasting exemplars of the active and inactive antimicrobial activity classes as shown in Figure 5 and in Figure 6. The contribution of the bitterness index (*bi*) is noticeably large while number of rotatable bonds (*nRotB*), molecular weight and polarizability are medium contributors. Only the (*nRotB*) and bitterness index change in the sign of their contribution while the rest are constant across the two comparisons. Overall, the importance of *bi* for class assignment is proven across the two examples.

**Figure 5:**
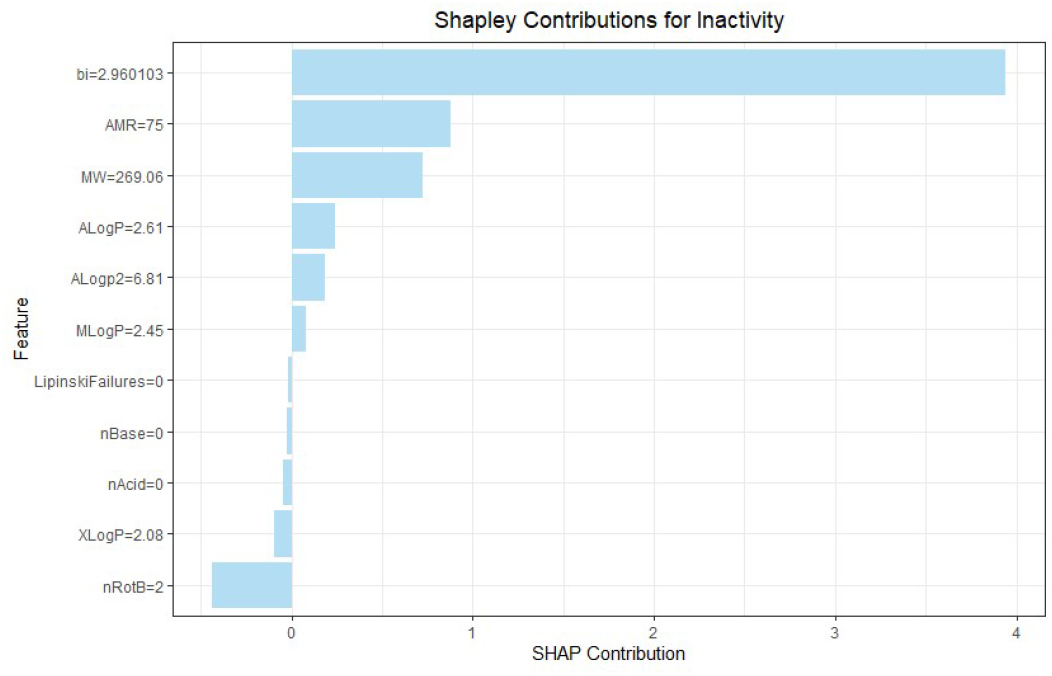
SHAP values applied to an inactive row showing the robust contribution of the *bi* relative other features.

**Figure 6:**
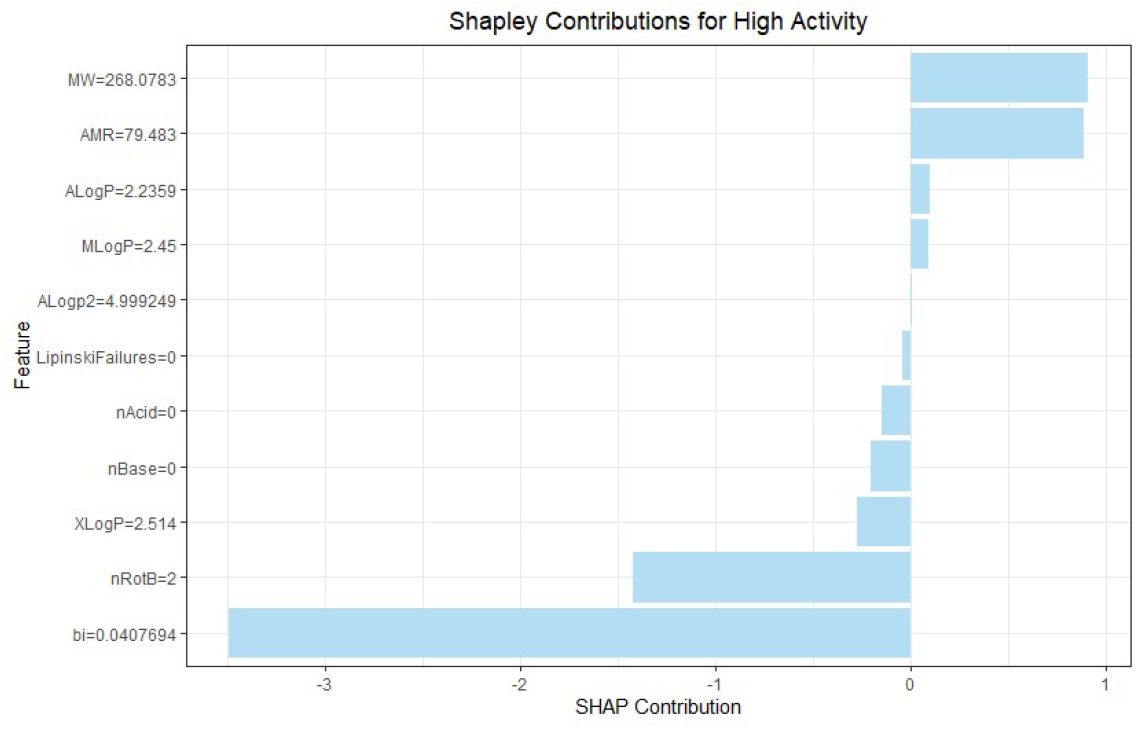
SHAP values applied to a highly active row showing the contribution of the *bi* relative other features. Directionality switch for bi between active and inactive rows demonstrates that bi is a distinguishing feature for antimicrobial activity

**Figure 7:**
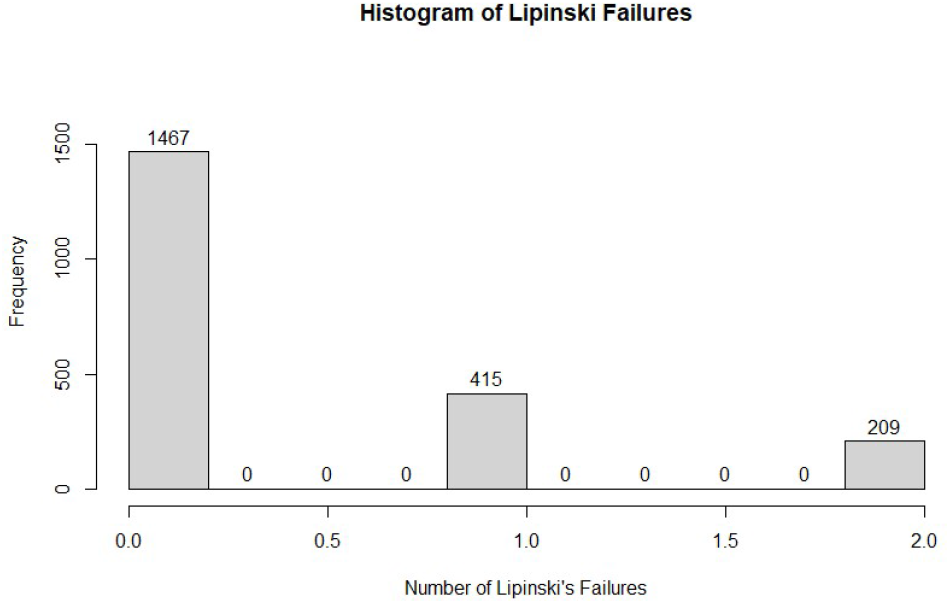
A histogram of the number of Lipinski’s failures shows that most have no failures but a significant number have either one or a couple of failures. This metric may be a basis for statistically matching the class members during training

**Figure 8:**
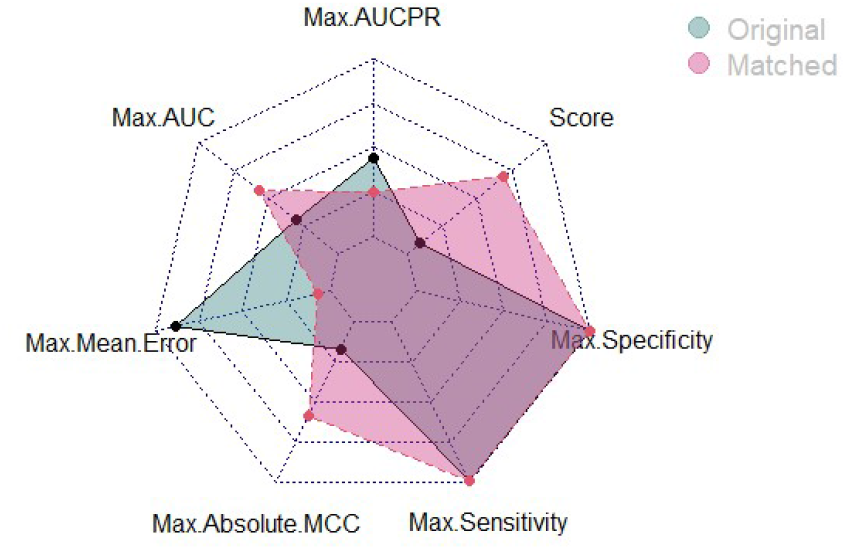
A radar plot showing that the matched model outperforms the traditional model especially in the MCC and mean error metrics. This is borne out by the overall score. Statistical matching enables fair comparisons and builds on the orthogonality in the predictor-predicted (category differentiation) set.

**Figure 9:**
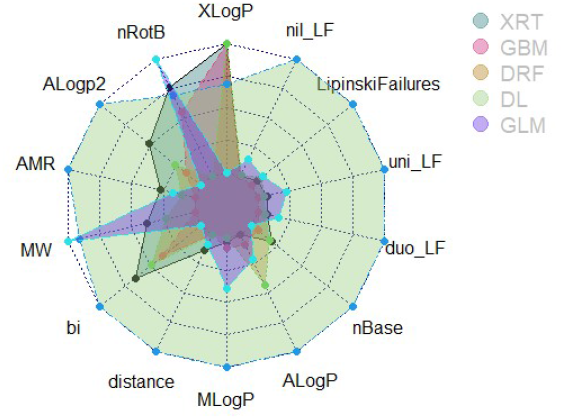
A radar plot showing the variable importance across the top performing model forms. In all cases, the value of the bitterness index is more muted but still a critical part of the modeling strength. Conversely, the partition coefficients have grown in relative importance. Lipinski Failures and associated encodings (*nil_LF, uni_LF, duo_LF*) are already known to be a relatively high scaled importance given their usefulness as de-confounding variables =. Note that the distance measures are extracted from the statistical matching process as a propensity score estimating the probability of being the interventional

#### 3.4.2 Comparison of modelling methods underscores high signal in the dataset compilation

By adapting the model form to include key driving factors, the number of viable algorithm choices may have been broadened. That means, predictive performance will remain relatively high/acceptable even as the algorithm type is altered. From Table IV, even though GBM and DRF are top ranked in 4 out of the 6 metrics, the XRT (extreme random forest trees) are not that far behind. This proves that the model choice made especially as to what factors are to be used as predictors, is a viable one.

**Table IV:**
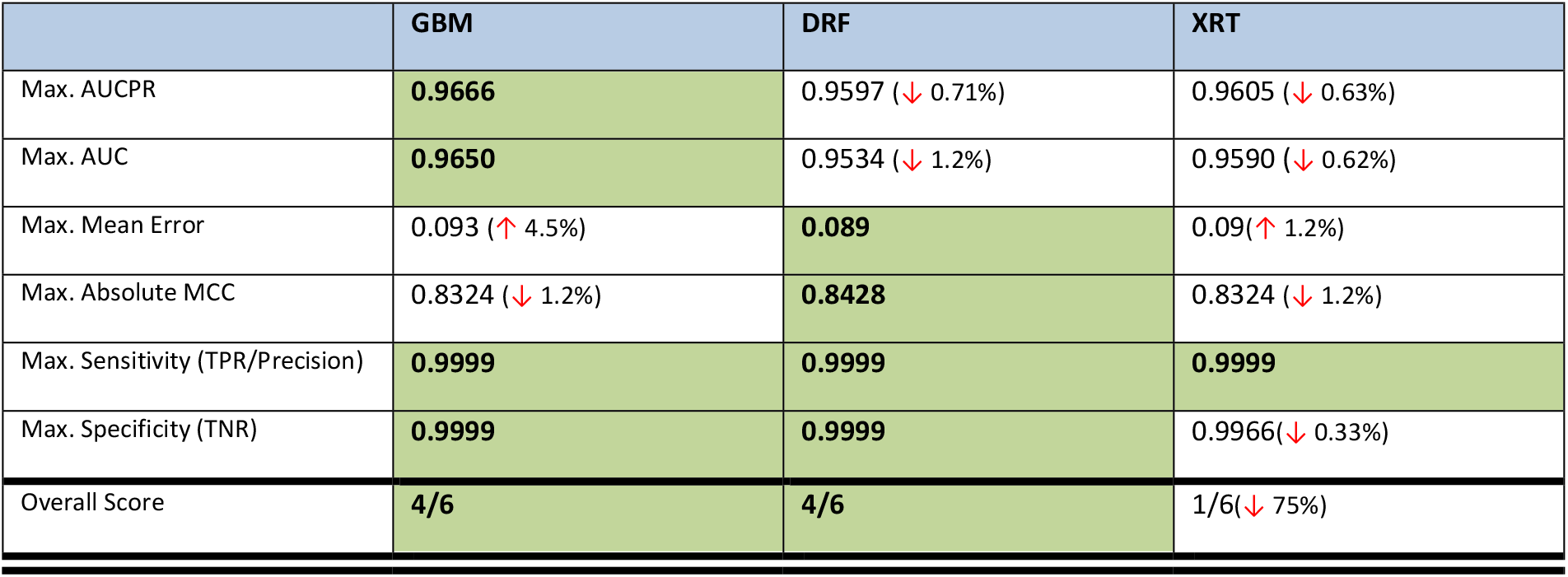
Performance comparison of the different algorithms demonstrates the data signal is robust and that the measurement choices regarding the response variable were sound in generating learnable contrasts between active and inactive antimicrobial activity classes. Absolute values are presented in each cell with the trends in adjacent brackets.

|  | GBM | DRF | XRT |
| --- | --- | --- | --- |
| Max. AUCPR | **0.9666** | 0.9597 (↓ 0.71%) | 0.9605 (↓ 0.63%) |
| Max. AUC | **0.9650** | 0.9534 (↓ 1.2%) | 0.9590 (↓ 0.62%) |
| Max. Mean Error | 0.093 (↑ 4.5%) | **0.089** | 0.09 (↑ 1.2%) |
| Max. Absolute MCC | 0.8324 (↓ 1.2%) | **0.8428** | 0.8324 (↓ 1.2%) |
| Max. Sensitivity (TPR/Precision) | **0.9999** | **0.9999** | **0.9999** |
| Max. Specificity (TNR) | **0.9999** | **0.9999** | 0.9966 (↓ 0.33%) |
| Overall Score | **4/6** | **4/6** | 1/6 (↓ 75%) |

**Table V:** Comparison of the statistically matched (SM) and traditional model with raw data (TM). The greatest change is observed with the improvement of the mean error & MCC. In contrast, the AUCPR dips negligibly relative the traditional (unmatched) model; other values are either the same or trending upwards.

|  | Traditional Model | Statistically Matched |
| --- | --- | --- |
| Max. AUCPR | **0.9666** | 0.9553(↓ 1.17%) |
| Max. AUC | **0.9650** | **0.9786**(↑ 1.41%) |
| Max. Mean Error | 0.093 | **0.0440**(↓ 52.69 %) |
| Max. Absolute MCC | 0.8324 | **0.9111**(↑ 9.45%) |
| Max. Sensitivity (TPR/Precision) | **0.9999** | **0.9999** |
| Max. Specificity (TNR) | **0.9999** | **0.9999** |
| Overall Score | **3/6**(↓ 40%) | **5/6** |

#### 3.4.3 Statistical matching with Lipinski’s Rules for de-confounded model learning

Potential confounding between the initial features and the generation of psychophysical parameters may lead to the statistical imbalance between classes and an inability to adequately parametrize each de-confounded relationship. A simple example may be that the partitioning constants are related to each other through an intermediary mid-line constant. Similarly, the psychophysical response is subject to intermediation by all constitutional parameters used as covariates in the model. Hence, it is a reasonable expectation that some confoundedness may converge upon a suboptimal model parametrization(19).

It is proposed that using bitterness as our chosen intermediating variable, one might identify fair categories of bitterness within the dataset. Based on these categories, a matched dataset displaying statistical balance on the covariates may better attribute the contributions to the model’s predictive power while increasing the diversity of model representations.

Having used a one-hot encoded mapping of Lipinski’s Failures to statistically match the data, the model was rebuilt with the results showing that the overall result was much improved with SM substantially improving the maximum absolute Matthew’s Correlation Coefficient (MCC, a metric balanced across all four confusion matrix quadrants). This demonstrates the value in improving data acquisition strategies across both the class-representation and the predictor-representation to enable fair comparisons.

A visual for the model changes shows that the statistically matched model fares better than the original unmatched model form with the main highlights being the lower mean error and the higher MCC. The overall score (“*Score*”) remains high as shown in

Using the matched model, we can re-examine the variable importance which still has the bitterness index playing a major role across the different model forms; however, there is a resurgence in the partitioning coefficients which matches general rules of thumb for drug discovery(20).

## 4 Conclusion

In conclusion, using a psychophysical index of bitterness (rendered by the TAS2R response to candidate molecules) alongside mainline physico-chemical descriptors to predict antimicrobial activity is methodologically sound. This work demonstrates that psychophysical indices may help to score and to identify relevant chemical spaces for molecular design and engineering. Further developments may examine safety and toxicity property predictions to accompany antimicrobial predictions. Adjacencies may exist to expand chemical adjuvation to reverse antimicrobial resistance as a means of extending the durability of legacy antimicrobials.

